# Analyses of cancer incidence and other morbidities in gamma irradiated B6CF1 mice

**DOI:** 10.1101/2020.03.26.009845

**Authors:** Alia Zander, Tatjana Paunesku, Gayle Woloschak

**Author notes:** Corresponding author: **Gayle Woloschak**.

## Abstract

With increasing medical radiation exposures, it is important to understand how different modes of delivery of ionizing radiation as well as total doses of exposure impact health outcomes. Our lab studied the risks associated with ionizing radiation by analyzing the Northwestern University Radiation Archive for animals (NURA). NURA contains detailed data from a series of 10 individual neutron and gamma irradiation experiments conducted on over 50,000 mice. Rigorous statistical testing on control mice from all Janus experiments enabled us to select studies that could be compared to one another and uncover unexpected differences among the controls as well as experimental animals. For controls, mice sham irradiated with 300 fractions died significantly earlier than those with fewer sham fractions. Using the integrated dataset of gamma irradiated and control mice, we found that fractionation significantly decreased the death hazard for animals dying of lymphomas, tumors, non-tumors, and unknown causes. Gender differences in frequencies of causes of death were identified irrespective of irradiation and dose fractionation, with female mice being at a greater risk for all causes of death, except for lung tumors. Irradiated and control male mice were at a significantly greater risk for lung tumors, the opposite from observations noted in humans. Additionally, we discovered that lymphoma deaths can occur quickly after exposures to high doses of gamma rays. This study systematically cross-compared outcomes of different modes of fractionation across a wide span of total doses to demonstrate that protraction modulated survival and disease status differently based on the total dose, cause of death, and sex of an animal. This novel method for analyzing the Janus datasets will lead to insightful new mechanistic hypotheses and research in the fields of radiation biology and protection.

## INTRODUCTION

Ionizing radiation is an unavoidable risk in daily life and understanding its biological impacts is important for setting radiation protection standards. Approximately half of humankind’s cumulative annual radiation exposure comes from natural sources, such as cosmic radiation and soil; the other half is derived from human-made sources including medical procedures and nuclear medicine (1). Most of the general population receives low dose chronic ionizing radiation exposures, accumulating to a few hundred mSv over a lifetime (2).

Calculating the risks associated with these chronic exposures is challenging because of the high background levels for the same diseases ionizing radiation can induce and the low overall effect of these lower dose/dose-rate exposures. There are several helpful data sources that researchers have utilized to help quantify these risks including radiation therapy studies, atomic bomb survivor data, and other epidemiological studies from workers in the field and nuclear disasters. The radiation doses given to patients for radiation therapy are much larger than standard exposure and are only used on a small segment of a patient, not whole-body exposures. The major source of data on whole body human exposures to gamma radiation is the Life Span Study (LSS) cohort that includes over 120,000 survivors of the atomic bombing in 1945 (3–6). While these data have been a remarkable resource for epidemiological studies determining risks associated with acute exposures (5, 7–13), extrapolation of health risks to humans exposed intermittently to lower doses of radiation remains uncertain. Different mathematical modelling approaches were used over the past 50 years to extrapolate health risks but they were met with variable enthusiasm from the scientific community (1, 14–18). Ultimately, epidemiological studies are affected by confounding factors and uncertainties making well-controlled animal studies a valuable resource to supplement conclusions from human studies (19).

We utilized the Northwestern University Radiation Archive for animals (NURA), a source of irradiated animal data documenting findings from Janus studies conducted between 1972 and 1989 at Argonne National Laboratory (ANL). Ten large volume experiments with B6CF1 mice were designed to determine the effects of acute and fractionated whole-body radiation on survival and causes of death (20–22). Over 50,000 male and female mice were exposed to acute or fractionated neutrons or gamma rays, at ages between 90-200 days or more than 500 days. Moribund animals were sacrificed and necropsy results were recorded. This experiment was one of the of the largest ever conducted in the USA; at the conclusion of these studies the Janus irradiator and other irradiation facilities at ANL were dismantled making it unlikely that experiments of this scope will be repeated.

Similar studies on other stains of mice were conducted in Europe (23) and more recent work conducted at the Institute of Environmental Sciences (IES) in Japan explored chronic exposures in a closely related animal strain. Tanaka and others compared findings on 4,000 B6C3F1 mice of both genders irradiated 22h daily for approximately 400 days to low dose rate gamma rays accumulating total doses of 0, 20, 400, or 8000 mGy (24). As the experiments performed at IES are similar to Janus experiments with regard to number of animals and total doses, we performed a side by side comparison of cancer incidence to determine biological similarities of these findings.

We examined whether fractionation, age at which a mouse was first irradiated, and gender modulate the overall death hazard and risk and frequency for specific causes of death. Our approach included use of general Cox proportional hazards models, cumulative incidence function models, and cause specific hazards models (25–27). We found that the two approaches to represent and evaluate competing risks from the same data complement each other and improve insight into effects of gamma ray fractionation. While this work cannot be directly translated into recommendations for radiation protection policies, it brings to our attention the fact that it should be possible to standardize comparisons between different types of fractionated exposures and perhaps fractionated versus chronic radiation exposures across a large span of total doses. A single mathematical formula cannot be used universally for conversion between any two possible radiation exposure scenarios, but it is possible that the growth of machine learning and artificial intelligence techniques will permit us to craft realistic approaches to predict changes in health complication spectra from one irradiation exposure to another, possibly even among different species. As we prepare for this future, it is necessary to ensure that we preserve radiation data archives with as much granularity as possible. This study is a prime example of utilizing archives by analyzing data in a new light to further augment our understanding of radiation biology.

## METHODS

### Data Selection - NURA

Argonne National Laboratory conducted a series of 10 large scale ionizing radiation lifespan studies on rodents between 1972 and 1989. These studies are now part of the NURA archive housed by the Woloschak laboratory and posted on the web, allowing access to all who are interested in this dataset (14, 21, 22, 28). Records list individual mouse information with the type of radiation, total dose, dose rate, fractionation schedule, age first irradiated, age at death, cause of death and, in many cases, detailed pathology analyses. All animals received whole body external beam ionizing radiation from cobalt 60 gamma rays or neutrons (20–22, 29). Most control mice were sham irradiated – transported from their housing location to the room with the irradiator turned off. Background radiation levels in animal housing rooms were closely monitored. Mice listed in **S1 Table** were censored due to early exit from the study because of causes unrelated to the experimental plan.

To ensure that any significant changes among different groups of mice were due to modulations in radiation exposure and not due to changes in baseline survival, we filtered out groups of mice that showed statistically significant survival probability differences. **S2 Table** details groups of mice that exhibited sufficient survival variation in control animals to warrant removal of the specific data from our analysis. Breeder mice were not used as controls for any of these analyses because of their unique housing conditions. Only *Mus musculus* species B6CF1 strain mice were used for this work; different species such as *Peromyscus leucopus* (white-footed deer mouse) were excluded from this study because of the species to species differences in response to radiation (30). Similarly, mice treated with radioprotectors (31) were also removed from this study. As a result of data refinement, only two of the ten experiments were completely removed from this study.

The predicted model output graphs from Cox Proportional Hazards (PH) analyses of sham irradiated control mice are shown in **S1 Fig** along with parameter estimates and p-values. Each overall model was significant due to sex, but no other covariates were significant in their respective models. Additionally, Kaplan Meier (KM) curves (32) in **S1 Fig e-h** validate the proportional hazards assumption for our model. To further validate our model, we used robustness tests, making small modifications to each variable in our models, shown in **S3 Table** and described in more detail in the Supplementary methods.

### Survival Analysis

Kaplan-Meier (KM) curves were used for categorical univariate survival analysis using the survfit function in the survival package in R (33, 34). Cox proportional hazard (PH) models were used to analyze survival over time with multivariate models that included a mixture of categorical and quantitative predictor variables and interactions between variables (25). The main models used for Cox PH with sham irradiated mice and shown in **S1 Fig** are as follows:

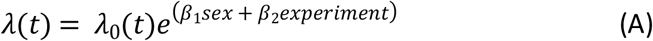

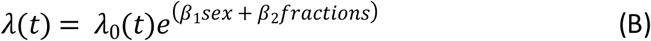

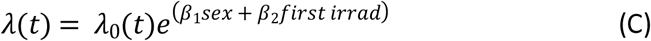

Our main Cox PH model for gamma irradiated mice is as follows:

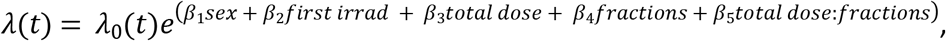

where *λ*(*t*) is the hazard function based on our set of covariates including sex, age first irradiated, total dose, number of fractions, and the interaction between total dose and fractions; β is a vector of their corresponding coefficients, and *λ*_0_(*t*) is the baseline hazard. All Cox PH models were performed using the coxph function from the survival package in R (34).

### Competing Risks Analysis

A competing risk is anything that decreases the likelihood of an outcome of interest. When looking at specific causes of death, all other causes of death fall into this category. For the competing risks analysis, we examined crude incidences, cause-specific hazard models, and cumulative incidence function (CIF) regression models (35). In the absence of competing risks, the cumulative incidence of events over time can be measured using one minus the Kaplan-Meier estimate of the survival function. In the presence of competing risks, the KM method results in upward biases for the CIF (26). We used the cuminc function from the cmprsk package in R to investigate crude, nonparametric incidences in the presence of competing risks (36).

For multi-variable regression analyses in the presence of competing risks, we used both cause specific hazards and CIF models. The cause specific hazards were estimated using the coxph function in R (34). All causes of death, excluding the event of interest, were censored. Concretely:

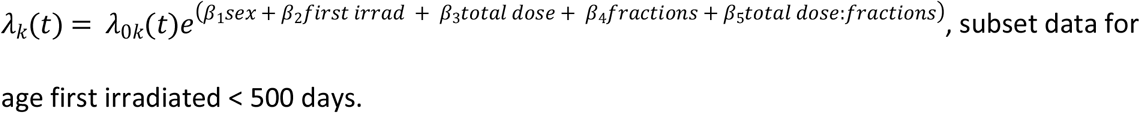

where *λ_k_*(*t*) is the hazard function for the k^th^ cause of death. The cause specific hazards method is used to determine the effect that covariates have on all event free subjects. The cumulative incidence function describes the overall probability of a particular outcome and does not depend on a subject being event free (26, 27, 37, 38). Concretely:

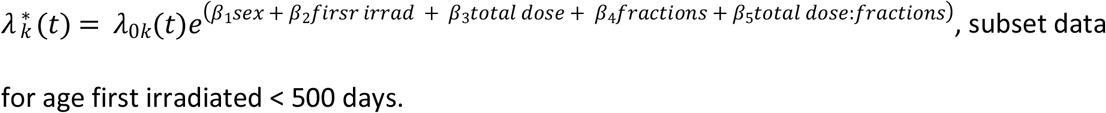

where 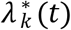 is the subdistribution hazard function for the k^th^ cause of death. Cumulative incidence hazards were estimated using the crr function in the cmprsk package in R (36).

### Cause of death groupings

We used data downloaded from the Janus website listed as “Grouped Macros,” which includes all pathologies found in animals at the time of death and categorizes them as lethal (L), contributory (C), or non-contributory (N). For the purposes of our investigation, we only examined lethal diseases. To make the data more robust for analyses, we grouped CODs into lymphomas, tumors other than lymphomas – referred to as tumors (sometimes separating them into lung tumors and tumors (no lung tumors)), non-tumors, or causes of death unknown (CDU). Specific analyses of diseases affecting the liver, lung, kidney, and vascular system for a subset of these data were previously conducted (39).

### Data reformatting for comparisons with IES data

Studies at IES involved low dose rate gamma irradiations of specific-pathogen free (SPF) B6C3F1 mice, F1 progeny of C57BL/6J females (B6) and C3H/HeJ males. While these mice are not identical to the B6CF1 mice (F1 progeny of C57BL/6J females (B6) and BALB/cJ males) used during the Janus experiments, they are matched in other respects. Both strains are F1 hybrids that share the same maternal strain C57BL/6.

During the IES studies, (SPF) B6C3F1 mice were irradiated with low dose rate ^137^Cs gamma rays for 22 hours a day, beginning the irradiations with acclimated 8-week old animals in a sterile environment. Chronic exposures of 0.05, 1.1 or 21 mGy/day continued for 400 days leading to total doses of 2, 40, or 800 cGy. Similar to the Janus experiments, many mice from the IES experiments were allowed to live out their entire lifespan, and each mouse was assigned a single cause of death (24, 40). These data were included in this study because of their similarity to the experiments carried out on B6CF1 mice during the NURA experiments.

To compare the Janus data included in this study with the results from the IES studies (24), we grouped Janus CODs to match IES CODs as closely as possible (**S4 Table**). The total doses used for Janus experiments spanned a larger range than those used for IES experiments. To make the comparisons more meaningful, we limited the Janus data used for this particular analysis to a subset of conditions that closely matched the IES dataset conditions. **S5 Table** provides information about the Janus data included in this comparison.

### Tools and Scripts

Files stored on github: https://github.com/aliazander

## Results

### Mice sham irradiated with 300 fractions had decreased survival

Control mice that received 300 fractions (5 fractions/week) of sham irradiation died significantly earlier than control animals that received fewer than 300 fractions of sham irradiation. This was evident in Cox PH models using sex and fractions as independent variables. This result held true with fractions treated as a continuous variable (**Fig 1A and S6 Table**) and as a categorical variable (**S2 Fig A and B**). KM curves showed a very similar trend to the predicted outcomes from the Cox PH models and validated the proportional hazards assumption of the Cox PH model (**S2 Fig C**). Because general stress is the only probable cause for the increased death hazard observed in this group of animals and irradiated animals exposed to 300 fractions most likely experienced the same stress, we excluded mice exposed to 300 fractions from our main analysis. These mice were included as part our robustness testing.

**Fig 1:**
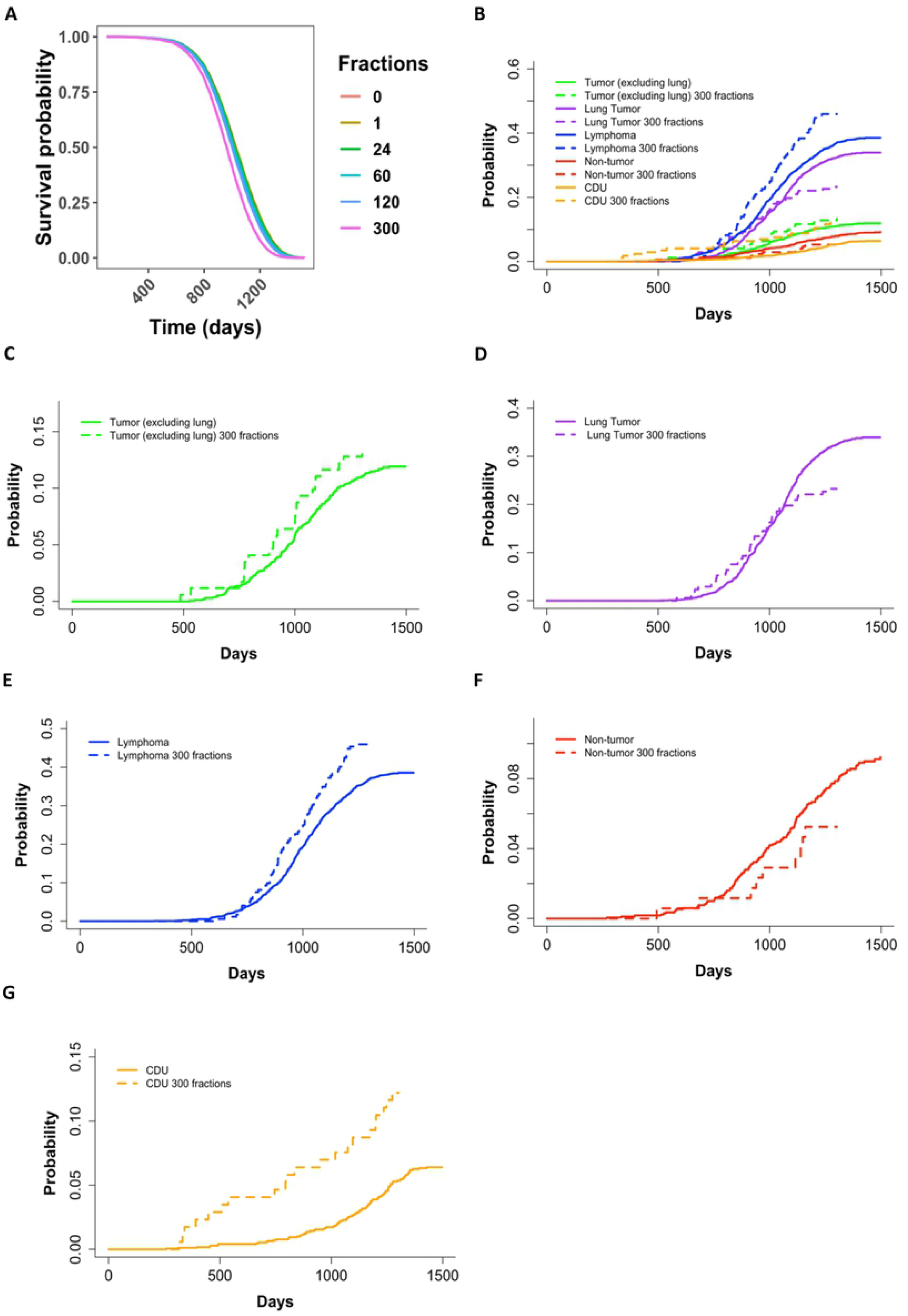
Survival probability output from Cox PH model for control mice with sex and the number of fractions as a continuous variable as independent variables (**A**). The predicted outcome shown in (**A**) are for male mice and output values shown in supplementary table 6. (**B**) Non-parametric cumulative incidence over time for specific causes of death grouped as tumors (excluding lung tumors), lung tumors, lymphomas, non-tumors, and CDUs. Dashed lines represent mice that received their sham irradiations in 300 fractions and solid lines represent mice that received their sham irradiations in fewer than 300 fractions. P-values for the differences of incidences between mice that received their sham irradiations in 300 fractions vs. mice that received their sham irradiations in fewer than 300 fractions: Tumor – 0.348, Lung Tumor – 0.038, Lymphoma - 0.017, Nontumor 0.222, CDU – 0.0002. For closer examination, we plotted each cause of death CIF individually – (**C**) tumors (excluding lung tumors), (**D**) lung tumors, (**E**) lymphomas, (**F**) non-tumors, and (**G**) CDUs.

### Mice sham irradiated with 300 fractions had significant changes in causes of death compared to other control mice

The decreased survival in mice that received 300 fractions compared to fewer than 300 fractions during their sham irradiations led us to investigate the specific causes of death for these two groups of mice. All animals sham irradiated with 300 fractions were male. Using non-parametric CIF, we found significant increases in lymphoma and CDU incidences and a significant decrease in lung cancer incidences in mice that received 300 sham irradiation fractions compared to all other sham irradiated male mice (**Fig 1**). Additionally, we examined how the number of fractions impacted survival probability over time through KM curves for each COD (**S2 Fig D-H**) and closer examination of the CIF curves in **fig 1C** by individually plotting each COD (**fig 1F-I**). The initial onset for lung tumor deaths (**fig 1F and S2 figE**) and CDU deaths (**fig 1I and S2 Fig H**) was earlier when mice received sham irradiations in 300 fractions, but there did not appear to be a difference for tumors (excluding lung tumors) (**fig 1E and S2 fig C**), lymphomas (**fig 1G and S2 fig E**), or non-tumors deaths (**fig 1H and S2 fig G**). The KM curves also supported these findings (**S2 fig C-H**).

### Aged mice were excluded from our analysis because of uneven experimental conditions

All control and gamma irradiated mice selected for this study (**S2 Table**) are represented in a box and whisker plot of age at death versus total dose with colors indicating the number of fractions used. Gamma irradiated mice received 21.57 to 4901 cGy (**Fig 2A**), with the maximal dose for acute exposures limited to 546 cGy. Because the LD_50/30_ for B6CF1 mice at 110 days of age is approximately 7 Gy (41), this maximum acute dose ensured that animals did not die from acute radiation syndromes. **Fig 2A** shows that fractionation had a larger impact on age at death as total doses increased. According to the Janus documentation, (21) the age first irradiated for mice was intended to be 100 days +/- 15 days, with a small subset of mice acutely irradiated at 500 days in order to investigate how age first irradiated impacted survival. Plotting the frequency of each age first irradiated, we found that the majority of mice were first irradiated within the expected range and there was a small group of mice that were first irradiated over 500 days old (**Fig 2C**). **Fig 2B** represents age at death against total dose with dark purple bars representing the aged mice and the light purple bars representing mice irradiated closer to 100 days of age. Due to the low sample size for aged mice, the large amount of leverage they would have on the overall model, the absence of sham irradiated aged mice, and the lack of direct dose comparisons with younger mice, we excluded these 560 mice from further analysis. This resulted in 11,618 total mice for the analysis on gamma irradiated mice.

**Fig 2:**
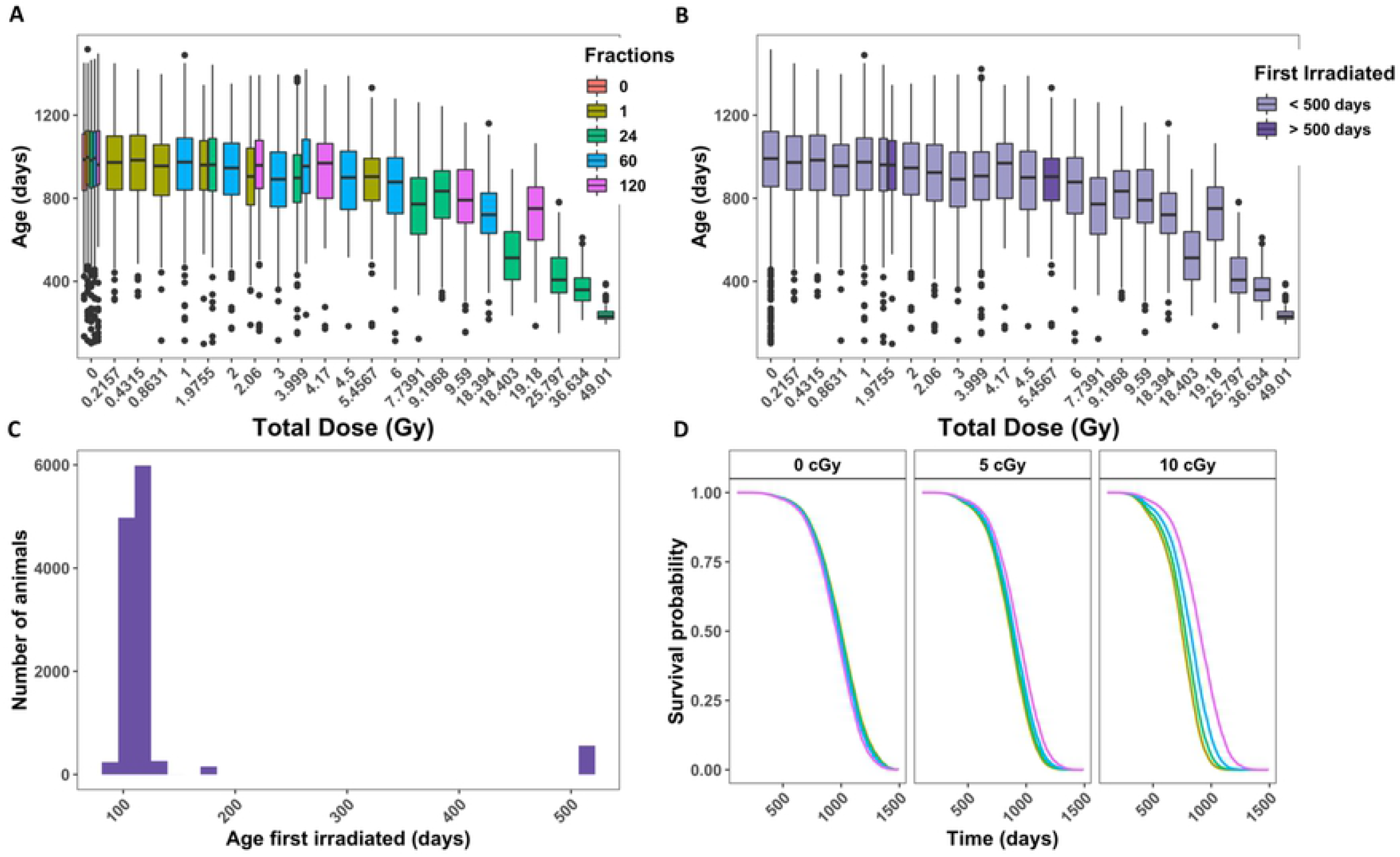
Analysis of animals filtered in Table 1 that were controls or gamma irradiated. (**A**) Box plot of age at death in days versus total dose in Gy. Colors indicate the number of fractions. (**B**) Histogram of the total number of animals versus age first irradiated in days. (**C**) Age at death in days versus total dose in Gy. Colors indicate whether a mouse was first irradiated before or after 500 days. (**D**) Representative graphs from Cox PH model output with age at death as the time scale and sex, age first irradiated, total dose, fractions, and the interaction between total dose and fractions as independent variables. The predicted outcomes shown are for female mice first irradiated at 120 days.

### Fractionation increased the overall survival probability in mice exposed to gamma rays

To determine how fractionation impacts survival, we used a Cox PH model with age at death as the time scale and sex, age first irradiated, total dose, fractions, and the interaction between total dose and fractions as independent variables. All independent variables were significant in the model, except for age first irradiated (**Table 1**). Given the small range of ages first irradiated included in our sample, this was an expected result. The main effect of fractions resulted in a positive coefficient from our model output, which corresponds to an increase in hazard. This result is the outcome of the difference in total doses of exposure for acute and fractionated radiation regimens. The maximum acute exposure was 5.4Gy, but the total doses for fractionated exposures reached as much as 49Gy, causing the interaction between total dose and the number of fractions to be the most relevant for determining the role of fractionation. The interaction term between fractions and total dose was highly significant and its interpretation is best understood by graphical representation of the model’s predicted outcome (**Fig 2D**). As the total dose increased, the beneficial effect of fractionation became more pronounced. This result was consistent throughout a series of robustness tests (**S3 Fig**). Additionally, we used KM curves to validate the proportional hazards assumption in our model and found parallel survival curves between groups based on sex, number of fractions, age first irradiated, and total dose (**S4 Fig**).

**Table 1:** Parameter estimates, hazard ratios with 95% confidence interval, and p-values for main Cox Proportional Hazards model in **Fig 1D**

| Variable | Estimate | Hazard Ratio (95% CI) | P-value |
| --- | --- | --- | --- |
| sexM | -0.17 | 0.84 (0.80, 0.89) | **<0.001** |
| Fractions | 0.002 | 1.00 (1.001, 1.003) | **0.001** |
| Total dose | 16.61 | 1.64E7 (8.2E6, 3.3E7) | **<0.001** |
| First irrad | -0.002 | 0.998 (0.996 1.00) | 0.137 |
| Fractions:Total dose | -0.109 | 0.89664 (0.88, 0.91) | **<0.001** |

### Fractionation significantly decreased the death hazard for mice dying from lymphomas, tumors, non-tumor, and causes of death unknown in gamma irradiated mice

When analyzing specific causes of death, it is important to consider the effects of competing risks. Cause specific hazards models are one type of competing risk model and their parameter estimates can be interpreted as the hazards for the specific event of interest. The cause specific hazards models for tumors, lymphomas, non-tumors, and CDUs all showed a similar trend (**Fig 3 A - D**) - as the dose increased, there was an increased rescue effect from fractionation, and more fractions corresponded to less hazard. Estimated model parameters showed that the interaction term between total dose and number of fractions was significant for all four categories of causes of death (**Table 2**). Sex was also significant in all models. Males had a higher death hazard for tumors, while females were at greater risk for all other causes of death, including lymphomas. Additionally, we examined the top two causes of death specifically - lung tumors and generalized non-thymic lymphomas (**S5 Fig**). Males were only at a greater risk of lung tumor death, while females had a higher hazard ratio for tumors excluding lung tumors and generalized non-thymic lymphomas.

**Fig 3:**
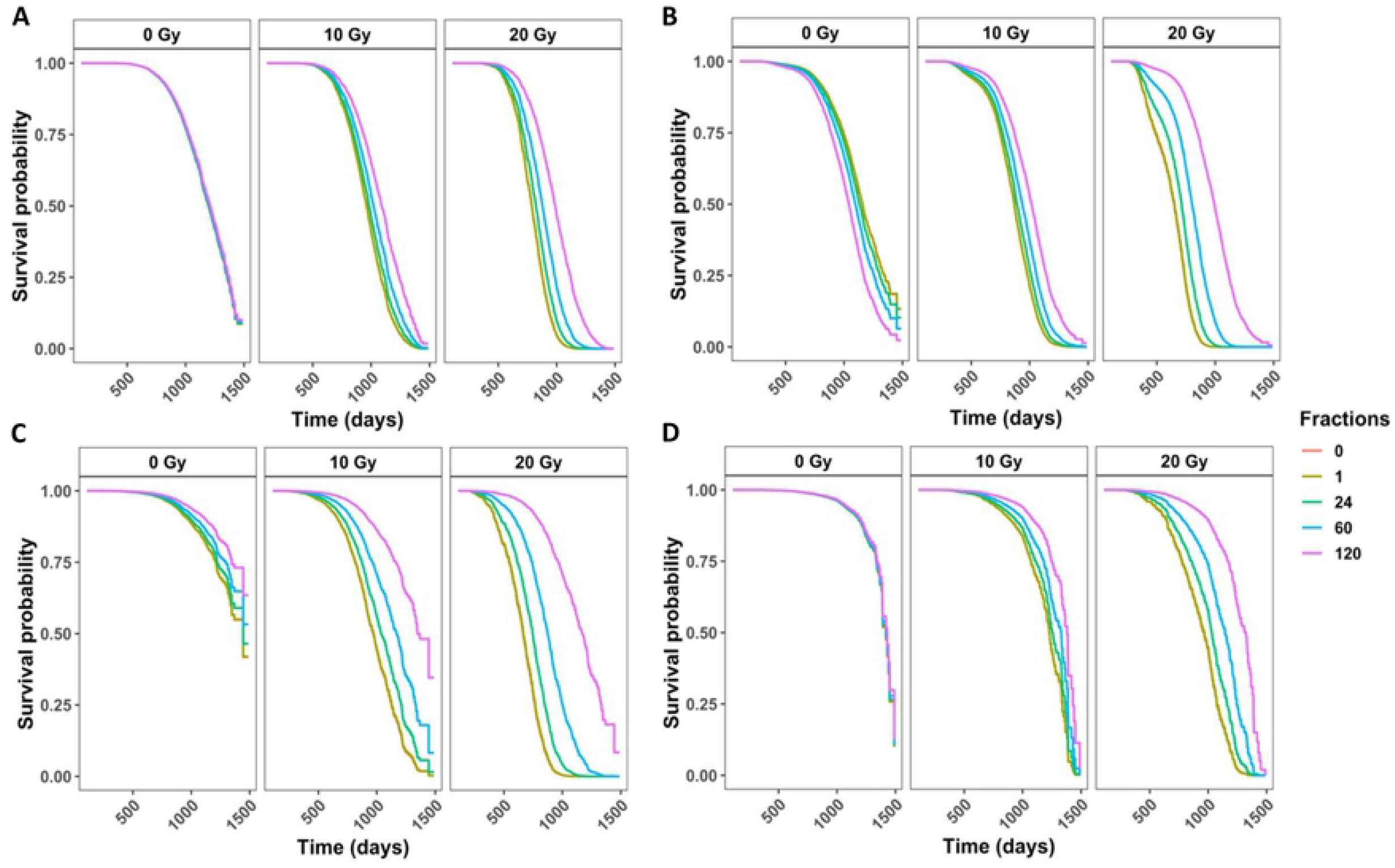
Competing risks models for specific causes of death in gamma irradiated mice with age as a time scale and sex, age first irradiated, total dose, fractions, and the interaction between total dose and fractions as independent variables. Survival curves for cause of death being (**A**) any solid tumors, (**B**) lymphomas, (**V**) non-tumors, and (**D**) cause of death unknown. Model estimates, confidence intervals and p-vaues are listed in the corresponding table 2. All four models have a significant interaction term between total dose and the number of fractions. The graphs represent predicted outcomes for female mice first irradiated at 120 days.

**Table 2:** Competing risks model output for cause specific hazards and subdistribution hazards

| COD | Independent Variable | Cause Specific Hazards |  |  | Subdistribution Hazards |  |  |
| --- | --- | --- | --- | --- | --- | --- | --- |
|  |  | Estimate | Hazard Ratio (95% CI) | P-value | Estimate | Hazard Ratio (95% CI) | P-value |
| Tumors | sexM | 0.195 | 1.21 (1.11, 1.33) | **<0.001** | 0.453 | 1.573 (1.446, 1.711) | **<0.001** |
|  | Fractions | -0.001 | 0.9994 (0.998, 1.001) | 0.535 | -0.003 | 0.997 (0.995, 0.999) | **0.002** |
|  | Total dose | 0.123 | 1.13 (1.11, 1.15) | **<0.001** | -0.063 | 0.939 (0.93, 0.948) | **<0.001** |
|  | First irradiation | -0.0001 | 0.9992 (0.996, 1.003) | 0.859 | 0 | 1 (1, 1) | 0.71 |
|  | Fractions: |  | 0.9994 (0.9991, | **0.001** |  |  |  |
|  | Total dose | -0.001 | 0.9998) |  | 0.001 | 1.001 (1, 1.001) | **<0.001** |
| Lymphoma | sexM | -0.37 | 0.69 (0.64, 0.75) | **<0.001** | -0.259 | 0.772 (0.72, 0.83) | **<0.001** |
|  | Fractions | 0.005 | 1.005 (1.004, 1.007) | **<0.001** | 0.004 | 1.004 (1.002, 1.005) | **<0.001** |
|  | Total dose | 0.166 | 1.18 (1.17, 1.19) | **<0.001** | 0.012 | 1.012 (1.004, 1.021) | **0.007** |
|  | First irradiation | -0.003 | 0.997 (0.993, 1.0001) | 0.058 | 0 | 1 (0.999, 1) | **.014** |
|  | Fractions: |  |  |  |  |  |  |
|  | Total dose | -0.001 | 0.999 (0.998, 0.999) | **<0.001** | 0 | 1 (1, 1) | .12 |
| Non-tumors | sexM | -0.50 | 0.61 (0.53, 0.69) | **<0.001** | -0.265 | 0.767 (0.676, 0.87) | **<0.001** |
|  | Fractions | -0.005 | 0.994 (0.991, 0.998) | **0.001** | -0.007 | 0.993 (0.99, 0.997) | **<0.001** |
|  | Total dose | 0.191 | 1.21 (1.19, 1.23) | **<0.001** | 0.07 | 1.073 (1.06, 1.086) | **<0.001** |
|  | First irradiation | -0.006 | 0.994, (0.988, 0.9999) | **0.009** | 0.001 | 1.001 (1.000, 1.001) | **<0.001** |
|  | Fractions: |  |  |  |  |  |  |
|  | Total dose | -0.001 | 0.999 (0.9990, 0.9995) | **<0.001** | 0 | 1 (1, 1) | .84 |
| CDU | sexM | -0.14 | 0.87 (0.71, 1.07) | 0.182 | -0.046 | 0.955 (0.79, 1.16) | 0.64 |
|  | Fractions | -0.001 | 0.999 (0.995, 1.003) | 0.667 | -0.002 | 0.998 (0.994, 1.002) | 0.34 |
|  | Total dose | 0.154 | 1.166 (1.14, 1.20) | **<0.001** | 0.002 | 1.002 (0.985, 1.02) | 0.82 |
|  | First irradiation | 0.001 | 1.001 (0.993, 1.01) | 0.782 | 3.96E-04 | 1.00 (1.00, 1.00) | 0.37 |
|  | Fractions: |  |  |  |  |  | 0.54 |
|  | Total dose | -0.001 | 0.999 (0.999, 0.999) | **0.011** | 0 | 1 (1, 1.001) |  |
| Lung tumors | sexM | 0.89 | 2.44 (2.12, 2.79) | **<0.001** | 1.19 | 3.30 (2.9, 3.8) | **<0.001** |
|  | Fractions | -0 | 0.9999 (0.997, 1.002) | 0.944 | -2.90E-03 | 0.997 (0.995, 0.999) | **0.0097** |
|  | Total dose | 0.11 | 1.12 (1.09, 1.14) | **<0.001** | -0.066 | 0.936 (0.925, 0.948) | **<0.001** |
|  | First irradiation | -0.004 | 0.996 (0.991, 1.001) | 0.130 | 0.001 | 1.001 (1, 1.001) | **.029** |
|  | Fractions: |  |  |  |  |  |  |
|  | Total dose | 0 | 0.9996 (0.999, 1.000) | 0.079 | 0.001 | 1.001 (1, 1.001) | **<0.001** |
| Tumors (excluding lung tumors) | sexM | -0.469 | 0.63 (0.55, 0.71) | **<0.001** | -0.38 | 0.684 (0.606, 0.773) | **<0.001** |
|  | Fractions | -0.001 | 0.999 (0.996, 1.001) | 0.348 | -0.003 | 0.997 (0.995, 1) | **0.021** |
|  | Total dose | 0.136 | 1.15 (1.12, 1.17) | **<0.001** | -0.057 | 0.944 (0.931, 0.958) | **<0.001** |
|  | First irradiation | 0.003 | 1.003 (0.998, 1.009) | 0.222 | -0.001 | 0.999 (0.998, 1) | **9.9E-03** |
|  | Fractions: |  |  |  |  |  |  |
|  | Total dose | -0.001 | 0.999 (0.9988, 0.9997) | **0.003** | 0.001 | 1.001 (1, 1.001) | **<0.001** |

### Cumulative incidence rates in deaths from lymphomas, tumors, non-tumors, and causes of death unknown in gamma irradiated mice varied greatly for each COD based on total dose and fractionation status

**Fig 4A** shows the non-parametric cumulative incidence of death for each of the main causes of death. Lymphomas were the most prevalent COD, followed by lung tumors, tumors (excluding lung tumors), non-tumors, and CDU. When we divided the data into control mice (**Fig 4B**) and gamma irradiated mice groups (**Fig 4C**), the first instances of death were observed around 750 days in control mice and around 250 days in gamma irradiated animals. These graphs were also subdivided by gender. For control and gamma irradiated mice, males had a lower incidence of lymphoma, tumors (excluding lung tumors), and non-tumors COD, while females had a much lower incidence of lung tumors. The differences between males and females were significant for both controls and gamma irradiated mice for all causes of death, except CDU (**Table 2**).

**Fig 4:**
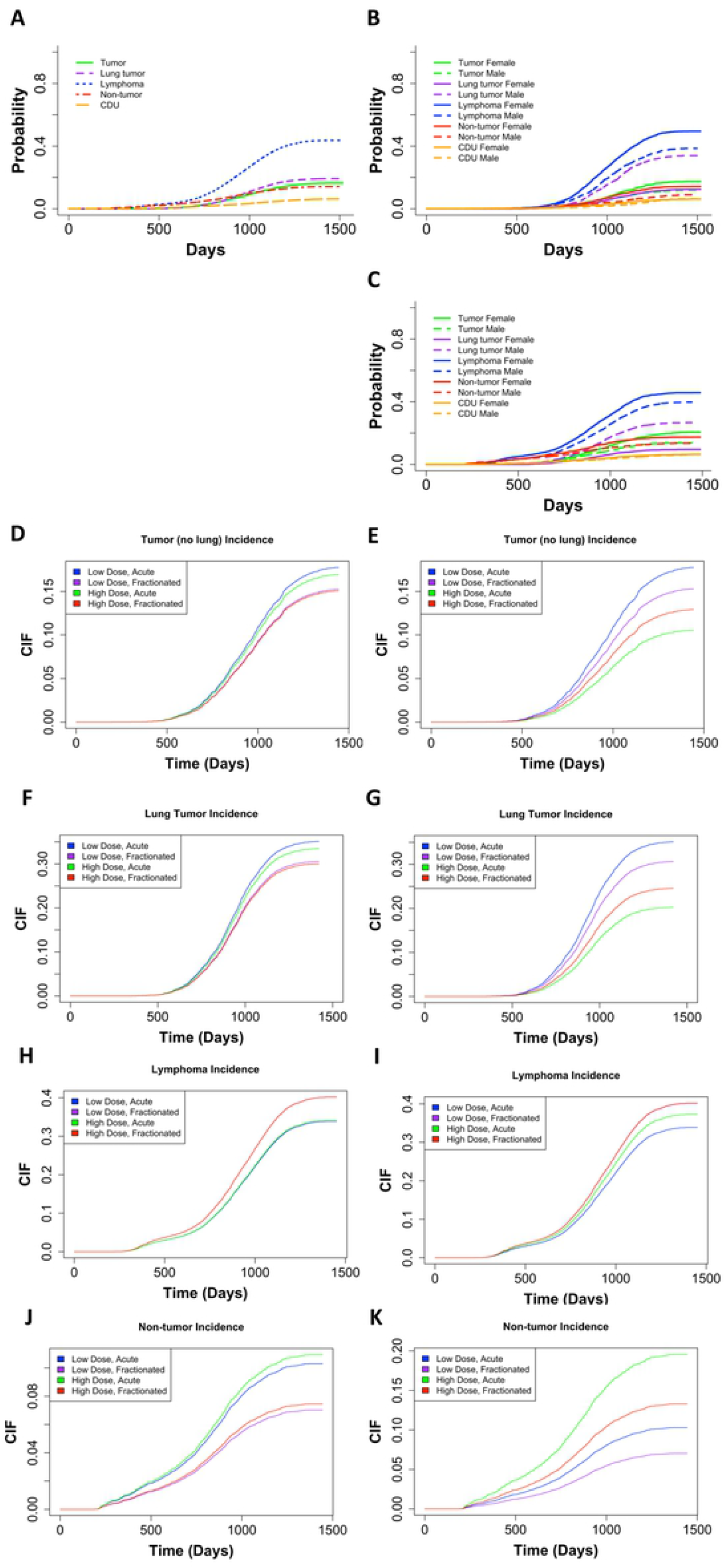
(**A**) Non-parametric CIF for the 5 main categories of COD without grouping. Non-parametric CIF for the 5 main categories of COD grouped my sex for control/sham irradiated mice (**B**) or gamma irradiated mice (**C**). P-values for differences in sex for each COD for mice graphed in (B): Tumor = 2.39E-06, lung tumor = 0, lymphoma = 1.99E-12, non-tumor = 5.04E-7, CDU 0.965, and for mice graphed in (C): Tumor = 3.97E-12, lung tumor = 0, lymphoma = 2.57E-08, non-tumor = 1.54E-05, CDU = 0.60. Predicted outcome under the following conditions: low dose = 0.1Gy, high dose = 1Gy for (**D**) tumors (excluding lung), (**F**) lung tumors, (**H**) lymphomas, and (**J**) non-tumors. Predicted outcome under the following conditions: low dose = 0.1Gy, high dose = 10Gy for (**E**) tumors (excluding lung), (**G**) lung tumors, (**I**) lymphomas, and (**K**) non-tumors. All predicted outputs are under the following conditions: Sex = males, acute = 1 fraction, fractionated = 60 fractions. Model output with parameter estimates, hazard ratios (95% confidence interval), and p-value are listed in **Table 2**.

In addition to calculating cause specific hazards for competing risks, it is important to examine subdistribution hazards, also known as cumulative incidence functions, using the Fine and Grey method (26, 27, 37, 38). The parameter estimates from subdistribution hazards have a less direct interpretation, but instead elucidate the overall probability of a particular outcome. We used the Fine and Grey method controlling for sex, age first irradiated, number of fractions, total dose, and the interaction between fractions and total dose, with competing risk groups as lymphoma, lung tumors, tumors (excluding lung tumors), non-tumors, and CDU. For tumors (excluding lung tumors) females were more susceptible but fractionation and increased dose both decreased tumor incidence (**Table 2).** By graphing predicted outcomes under varying conditions, we discovered that when the difference between high and low total doses is small (10cGy vs 100cGy), fractionation is the biggest determinate for tumor incidence, with acute exposures resulting in the most tumors (**Fig 4D**). Conversely, when the high dose was increased to 1000cGy, total dose became the dominant factor and low dose conditions resulted in the most tumor incidences (**Fig 4E**). Under all conditions, low dose acute exposures resulted in the greatest tumor incidence. Fractionation and dose had the same impact on lung tumor incidence as in all other tumor incidence (**Fig 4 F and G**). However, males were more likely than females to die of lung tumors specifically, which matches the cause specific hazards results (**Table 2**).

Examining lymphoma deaths, females were at a greater risk of death than males and increasing the total dose and number of fractions both increased the risk of death (**Table 2**). Predicted outcomes showed that with a 10-fold difference between high and low total doses, fractionation was the main determinate for lymphoma incidence and fractionated exposures resulted in the most lymphoma cases (**Fig 4H**). Predicted outcomes with a 100-fold difference between high and low total doses resulted in total dose became the dominant factor and high dose conditions produced the most lymphoma incidences (**Fig 4I**). For all conditions, high dose fractionated exposures resulted in the greatest lymphoma incidence, which is the exact opposite from the trend observed for tumor deaths.

Non-tumor deaths were more prevalent in female mice compared to male mice, total dose increased the probability of non-tumors, and fractionation decreased the probability of a non-tumor death (**Table 2**). When we examined the predicted outcome using a 10-fold difference between high and low total doses, fractionation had the greatest impact on lymphoma incidence, with acute exposures resulting in the most non-tumor cases (**Fig 4J**). We observed that dose became the dominant factor when we assessed a 100-fold difference between high and low total doses, and high dose conditions resulted in the most non-tumor deaths (**Fig 4K**). High dose acute exposures resulted in the greatest non-tumor incidence consistently for all conditions we analyzed.

### Lymphoma and non-tumors both had higher incidence rates with high doses at earlier times than previously anticipated

The CIF regression models for death by lymphomas and non-tumors both demonstrated a shoulder along the CIF curve around 250-500 days (**Fig 4 H-K**). When we filtered out total doses above 6Gy (**S6 Fig**), this shoulder disappeared for lymphoma deaths and non-tumor deaths. Because tumors (excluding lung tumors) and lung tumors (**Fig 4 D-G**) did not exhibit the same shoulder, these results act as an internal negative control. After filtering out mice exposed to total doses over 6Gy for tumors and lung tumors, the shape of the CIF curves did not change (**S6 Fig B, D**).

### Janus datasets showed similar results to IES data in male versus female comparisons and dose response trends for causes of death

Several large-scale chronic exposure studies were done at IES using both genders of B6C3F1 mice. Work by Tanaka and others (24) at IES was focused on specific causes of death in response to gamma irradiation and we compared these data with Janus data. As mentioned in the methods, the B6C3F1 mice strain used at IES was genetically similar to the B6C3F1 mice used in the Janus experiments. Both strains of mice came from crosses of the same radiosensitive strain of female mice (BALB/c) and relatively radioresistant male mice. When examining cause of death between groups (**Table 3**), we found that hematopoietic system diseases are the most common cause of death for both sets of mice. Additionally, females died of hematopoietic diseases more than males in both datasets. The next most common cause of death for Janus mice was respiratory disease, a result driven by the high incidence of lung cancer in B6CF1 mice from Janus studies. B6C3F1 mice died much less frequently of respiratory disease compared to the B6CF1 mice. However, respiratory diseases were more common in male mice in both datasets. Finally, in male B6C3F1 mice, digestive diseases were much more frequent than in animals used in Janus experiments.

**Table 3:**
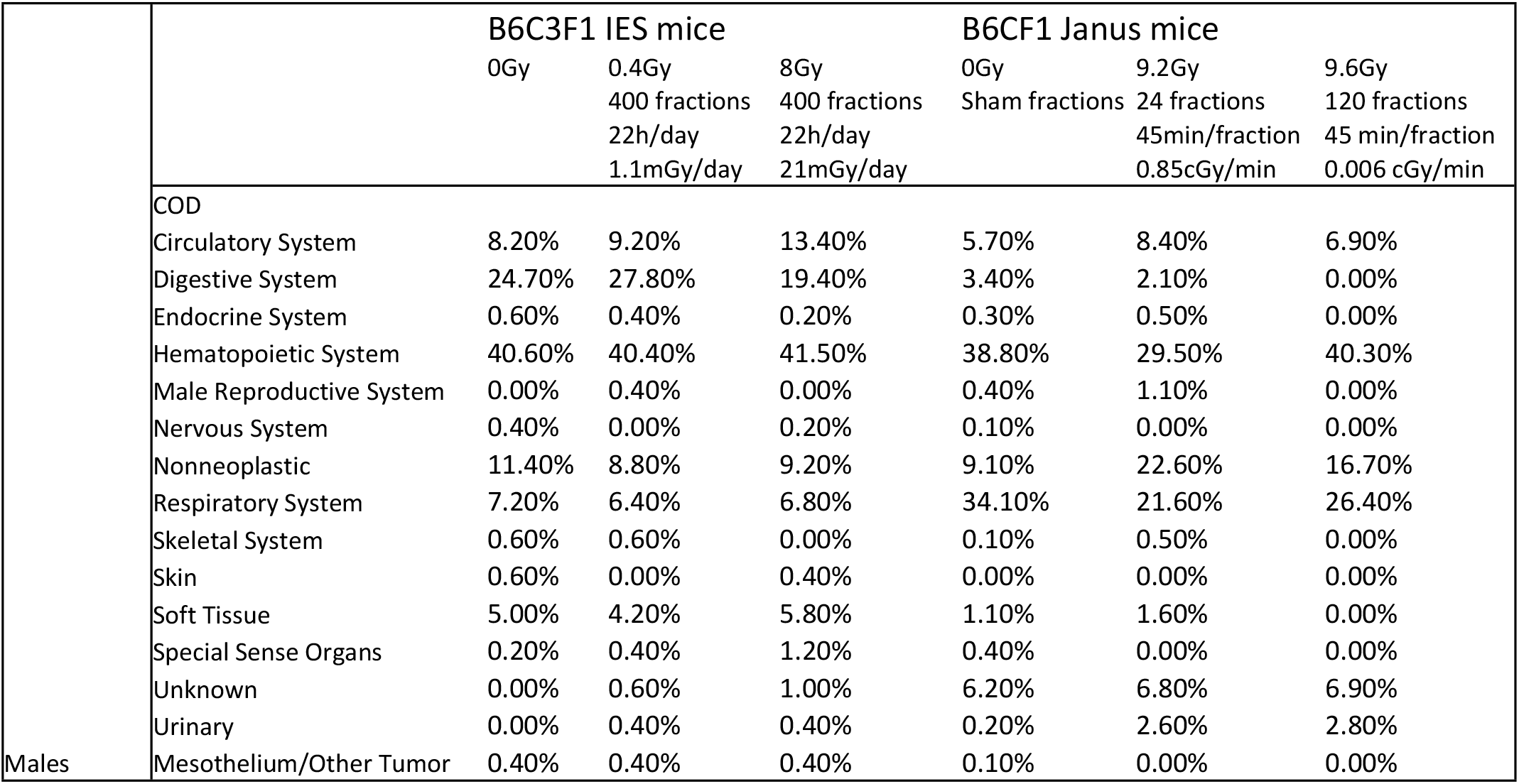
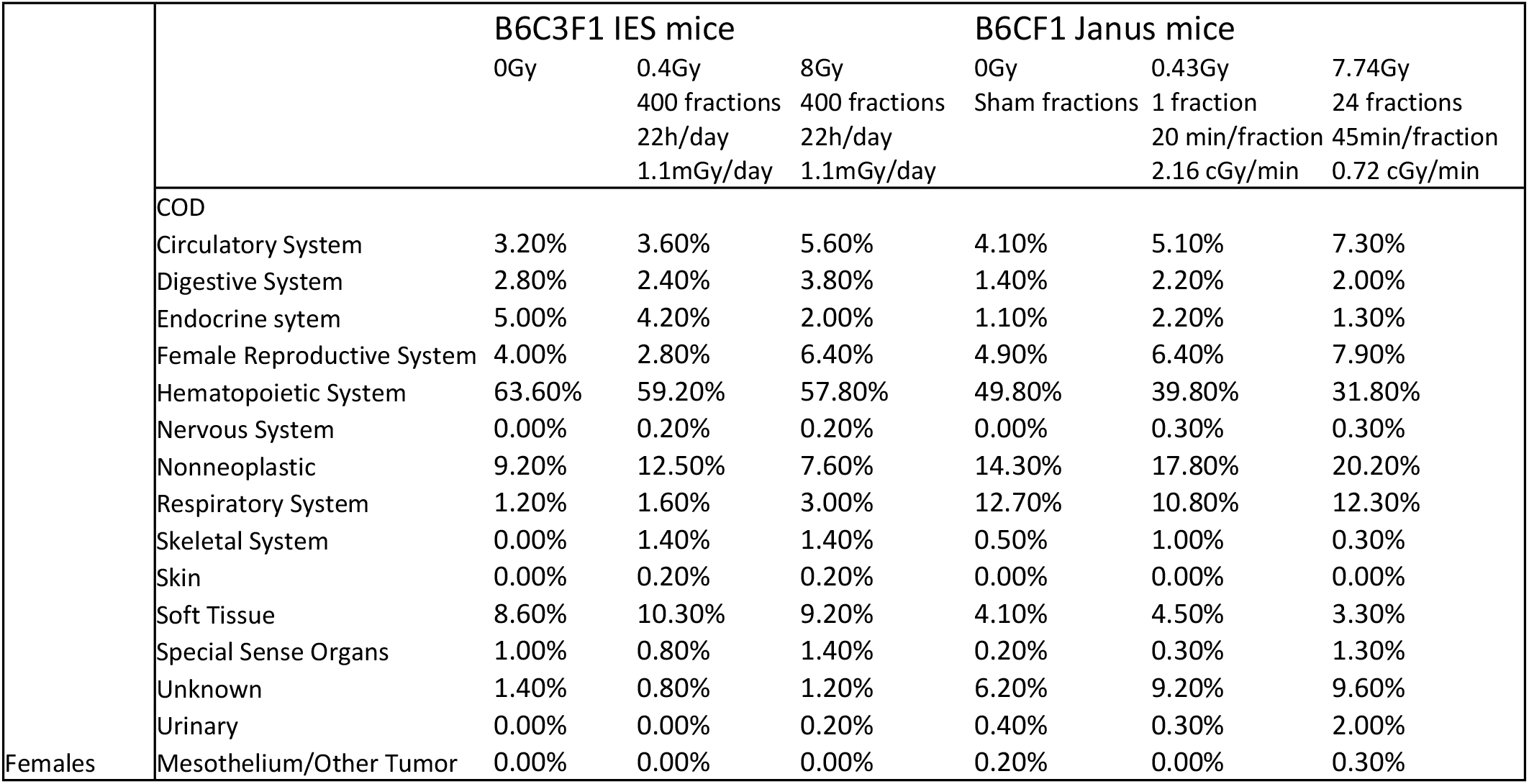
The percentage of deaths due to each individual cause of death listed for a comparison between B6CF1 Janus mice and B6C3F1 IES mice.

## Discussion

Janus experiments were analyzed in many different ways over the years (14, 17, 20, 30, 31, 39, 42–47) and each new approach for analysis of these data brought novel information about the effects of dose fractionation. By analyzing control mice in a way that allowed us to pool Janus experiments together, we gained statistical power to run tests on the importance of fractionation for specific causes of death. The Janus experiments were originally designed with this in mind and taking advantage of the consistency between experiments for a large-scale study was extremely effective. We were able to determine under which circumstances fractionation had a rescuing effect and track changes in risk based on gender for specific causes of death. Our method for pooling data together can be used for future analysis on the Janus dataset using different modeling techniques and answering novel biological questions.

One of the most interesting findings from the control mice analysis was that mice sham irradiated with 300 fractions died significantly earlier than animals exposed to fewer fractions during their sham irradiations. Mice given 300 sham irradiation fractions died earlier due to tumors and CDU. The simplest and most likely explanation for this phenomenon is general stress caused by frequent exposure to unfamiliar circumstances. Sham exposures involved transporting the mice from the room where they were housed to the room with the irradiator. The irradiator was turned off and there was no excess radiation in the room. It is known that transporting mice induces a stress response (48, 49). Further investigation of this mouse cohort may provide us with new insights into the stress response. Analyzing available tissue samples from these mice could enable us to explore cytological or molecular indicators of stress. These results would not only be beneficial for animal studies in radiation biology, but for any investigators utilizing animals in their research.

In all comparisons between acute and fractionated exposures, fractionation significantly decreased the death hazard in gamma irradiated mice. Moreover, fractionation was equally protective for all four pooled categories of diseases (lymphoma, tumors, non-tumors, and CDU) and specific diseases such as non-thymic lymphoma. Lung tumors were the only causes of death that were not significantly affected by fractionation. B6CF1 males were at a significantly higher risk for lung tumors than females. Interestingly, when we excluded lung tumors and examined all other types of tumors, males were at a lower risk than females for all other causes of death. Heidenreich et al. investigated lung tumors from the Janus datasets using Kaplan-Meier plots and the two-step clonal expansion (TSCE) model (50). They concluded that males had a higher lung cancer risk than females in control and gamma irradiated mice. They also found that more fractions administered over a longer duration resulted in less lung tumor risk, again agreeing with the results we found using Cox PH.

Examining CIFs to determine the probabilities for death due to distinct diseases under varying conditions produced many intriguing results. Increasing the number of fractions and increasing the total dose both decreased the incidences of tumors and lung tumors specifically. This finding is not surprising because tumors in mice develop more slowly than lymphomas and even more slowly than non-tumors such as radiation induced pneumonitis. Therefore, a mouse exposed to a high dose of gamma rays would most likely die of non-cancer cause of death before a tumor has time to fully advance.

Mice exposed to fractionated irradiation died from lymphoma more frequently when compared to mice that received acute exposures, because mice exposed to acute exposures developed more non-tumors. Non-tumors were the most common in response to higher doses and acute exposures.

Lymphomas are easily induced in response to ionizing radiation in rodents. It is typically considered a risk associated with lower doses of ionizing radiation compared to non-tumors. However, our results showed an early increase in lymphoma incidences when the dose delivered to animals was above 6 Gy. A 6Gy cutoff was chosen because all of the animals exposed to doses above 6 Gy received fractionated irradiation. This was done because the LD_50/30_ dose for B6CF1 mice is 6.54 or 6.75 Gy for males and females respectively (25) and acute exposures above 6Gy would result in an animal death rate incompatible with robust experimental data. In mice that received total doses over 6Gy (all fractionated exposures), lymphoma deaths began as early as 300 days. The early death shoulder observed in CIF curves for lymphomas is no longer present when excluding data for total doses above 6Gy. Non-tumor deaths demonstrated the same shoulder when the full range of total doses were included to fit the model. Again, the shoulder disappears after removing the data for total doses over 6Gy.

We compared the Janus dataset on fractionated radiation with the IES datasets on chronic irradiation. This comparison between B6CF1 mice used in Janus experiments and B6C3F1 mice used by Tanaka and others (24) showed a high degree of similarity despite different radiation delivery approaches and genetic differences between the two strains. The most pronounced differences between B6CF1 and B6C3F1 mice were associated with male mice, which could be attributed to these two hybrid strains differing by the paternally contributing mice. While respiratory system complications affected both F1 mouse hybrids discussed here, not all mice have the same association between gender and radiation associated respiratory diseases. For example, RFM mice exposed to x-rays males were at less risk than females for lung tumors, indicating that differential gender susceptibilities to lung tumors are strain specific (51–53). IES results also showed more death due to digestive diseases, with the effect being most obvious in male mice. Janus mice were not kept in a sterile environment nor fed sterile food. The bacteria present in the guts of Janus mice would have increased the local immune response and may explain the lower percentage of deaths due to digestive diseases. These noticeable differences in digestive and respiratory disease proportions could be explained by several differences in housing. Not only were IES mice specific pathogen free, while Janus mice were not, but it is also likely that standard housing conditions changed between 1972 and 2004. Beyond standard conditions over time, conditions are likely variable across universities and countries, as well. In humans, females have been shown to be at a greater risk of lung cancer than males (54, 55). Determining the cause for changes in lung tumor sensitivity in response to ionizing radiation between male and female mice could lead to a better understanding of the radiation induction of lung tumors. Given our current, limited amount of information, RFM mice appear to be a better model system for simulating gender differences of humans for lung tumor risk.

In conclusion, we propose to continue to evaluate the NURA database using different models and applying them to different subsets of data in order to outline the finer nuances of consequences of radiation exposures. The differences we described in radiation exposure outcomes that change with alterations in radiation delivery highlight that biological responses to whole body irradiation most likely cannot be described by a single factor that could be applied for the entire spectrum of possible fractionation scenarios.

## Acknowledgements

The authors would like to thank Edward Malthouse for his input on the statistical methods used and Carissa Ritner and Benjamin Haley for their constant support and thoughtful discussions.

## Supplemental Information

**S1 Table:** Cause of Death (COD) codes for censored mice and the number of mice in each group.

**S2 Table:** Description of which mice were removed from our analysis, the corresponding reasoning, and the resulting sample size. N=4330 for control mice and N=7848 for gamma irradiated mice.

**S1 Fig:** Cox proportional hazard (PH) survival curves produced using all data filtered by the criteria in Table 1 that also had a total dose of 0Gy. Age at death was used as a time scale with sex and (A) experiment, (B) number of fractions, or (C) the age first irradiated as independent variables. The predicted outcomes shown are for female mice. Parameter estimates and p-values from each model are shown in D. Kaplan Meier curves with (E) sex, (F) number of fractions, (G) age first irradiated, and (H) experiment group were evaluated to test the proportional hazards assumption of these models.

**S3 Table:** Robustness tests for Cox Proportional Hazards models for controls analysis.

**S4 Table:** Grouping B6CF1 Janus cause of death codes to match B6C3F1 IES categories

**S5 Table:** Janus experiments used for comparisons with IES chronic irradiation data

**Supplementary Table S6:** Survival probability output from Cox PH model with sex and the number of fractions as a continuous variable in control mice. Corresponding graph in figure 1A.

**S2 Fig:** Survival probability output from Cox PH model with sex and the number of fractions as a categorical variable control mice (A/B). The predicted outcomes shown (A) are for male mice. Kaplan Meier curve for control mice showing differences in survival probability based on the number of fractions used during sham irradiation treatment (C). Kaplan Meier curves showing how the number of fractions impacts survival in control male mice that died of solid tumors (excluding lung tumors) (D), lung tumors (E), lymphomas (F), non-tumor diseases (G), and CDU (H).

**S3 Fig:** Robustness testing for gamma irradiation main model. CoxPH model output from the base model with the following changes: (A/B) including mice irradiated at all ages, (C/D) stratifying by sex, (E/F) treating fractions as a categorical variable, (G/H) including exact fraction number and age first irradiated for control samples, (I/J) including mice with total dose received in 300 fractions, and (K/L) including mice with total dose received in 300 fractions and fractions treated as a categorical variable. The graphs represent predicted outcomes for female mice first irradiated at 120 days.

**S4 Fig:** Kaplan Meier curves showing survival probably vs. time in gamma irradiated and control mice after the filtering shown in Table 1 and removing mice first irradiated after 500 days. KM curves compare (A) sex, (B) number of fractions, (C) age first irradiated, and (D) total dose.

**S5 Fig:** Competing risks models for specific causes of death in gamma irradiated mice with age as a time scale and sex, age first irradiated, total dose, fractions, and the interaction between total dose and fractions as independent variables. Survival curves for cause of death being (A) lung tumors, (B) all tumors (excluding lung tumors), and (C) non-thymic lymphomas. The corresponding model output with parameter estimates, hazard ratios with a 95% confidence interval, and p-values are listed in table 2 for lung tumors and all tumors (excluding lung tumors). (D) Model output for non-thymic lymphoma. The graphs represent predicted outcomes for female mice first irradiated at 120 days.

**S6 Fig:** CIF regression model output using a total dose cutoff of 6Gy for tumor (no lung) (A), lung tumor (C), lymphoma (E), and non-tumor deaths (G). Predicted output from CIF model for tumor (no lung) (B), lung tumor (D), lymphoma (F), and non-tumor deaths (H). All predicted outputs represent males first irradiated at 120 days under the following conditions: low dose = 0.1Gy, high dose = 6Gy, acute = 1 fraction, fractionated = 60 fractions.

## References

1. Measurements NCoRPa. Ionizing radiation exposure of the population of the United States. National Council on Radiation Protection and Measurments; 2009. Report No.: 160.

2. Mettler FA, Jr., Huda W, Yoshizumi TT, Mahesh M. Effective doses in radiology and diagnostic nuclear medicine: a catalog. Radiology. 2008;248(1):254–63.

3. Grant EJ, Brenner A, Sugiyama H, Sakata R, Sadakane A, Utada M, et al. Solid Cancer Incidence among the Life Span Study of Atomic Bomb Survivors: 1958-2009. Radiation research. 2017;187(5):513–37.

4. Hsu WL, Preston DL, Soda M, Sugiyama H, Funamoto S, Kodama K, et al. The incidence of leukemia, lymphoma and multiple myeloma among atomic bomb survivors: 1950-2001. Radiation research. 2013;179(3):361–82.

5. Ozasa K, Cullings HM, Ohishi W, Hida A, Grant EJ. Epidemiological studies of atomic bomb radiation at the Radiation Effects Research Foundation. Int J Radiat Biol. 2019;95(7):879–91.

6. Ozasa K, Grant EJ, Kodama K. Japanese Legacy Cohorts: The Life Span Study Atomic Bomb Survivor Cohort and Survivors’ Offspring. J Epidemiol. 2018;28(4):162–9.

7. Brenner AV, Preston DL, Sakata R, Sugiyama H, de Gonzalez AB, French B, et al. Incidence of Breast Cancer in the Life Span Study of Atomic Bomb Survivors: 1958-2009. Radiation research. 2018;190(4):433–44.

8. Sadakane A, French B, Brenner AV, Preston DL, Sugiyama H, Grant EJ, et al. Radiation and Risk of Liver, Biliary Tract, and Pancreatic Cancers among Atomic Bomb Survivors in Hiroshima and Nagasaki: 1958-2009. Radiation research. 2019.

9. Sakata R, Preston DL, Brenner AV, Sugiyama H, Grant EJ, Rajaraman P, et al. Radiation-Related Risk of Cancers of the Upper Digestive Tract among Japanese Atomic Bomb Survivors. Radiation research. 2019.

10. Sentani K, Ogawa I, Ozasa K, Sadakane A, Utada M, Tsuya T, et al. Characteristics of 5015 Salivary Gland Neoplasms Registered in the Hiroshima Tumor Tissue Registry over a Period of 39 Years. J Clin Med. 2019;8(5).

11. Sugiyama H, Misumi M, Brenner A, Grant EJ, Sakata R, Sadakane A, et al. Radiation risk of incident colorectal cancer by anatomical site among atomic bomb survivors: 1958-2009. Int J Cancer. 2019.

12. Sugiyama H, Omonishi K, Yonehara S, Ozasa K, Kajihara H, Tsuya T, et al. Characteristics of Benign and Malignant Bone Tumors Registered in the Hiroshima Tumor Tissue Registry, 1973-2012. JB JS Open Access. 2018;3(2):e0064.

13. Utada M, Brenner AV, Preston DL, Cologne JB, Sakata R, Sugiyama H, et al. Radiation Risks of Uterine Cancer in Atomic Bomb Survivors: 1958-2009. JNCI Cancer Spectr. 2018;2(4):pky081.

14. Tran V, Little MP. Dose and dose rate extrapolation factors for malignant and non-malignant health endpoints after exposure to gamma and neutron radiation. Radiat Environ Biophys. 2017;56(4):299–328.

15. Ruhm W, Azizova TV, Bouffler SD, Little MP, Shore RE, Walsh L, et al. Dose-rate effects in radiation biology and radiation protection. Ann ICRP. 2016;45(1_suppl):262–79.

16. SSK. Dose-and dose-rate-effectiveness factor (DDREF), Recommendation by the German Commission on Radiological Protection with scientific grounds. 2014.

17. Haley BM, Paunesku T, Grdina DJ, Woloschak GE. The Increase in Animal Mortality Risk following Exposure to Sparsely Ionizing Radiation Is Not Linear Quadratic with Dose. PLoS One. 2015;10(12):e0140989.

18. Academies. BoRERDoEaLSNRCotN. Health Risks from Exposure to Low Levels of Ionizing Radiation: BEIR VII Phase 2. 2006.

19. Radiation UNSCotEoA. UNSCEAR 2012 Report Annex B. Uncertainties in risk estimates for radiation-induced cancer.; 2012.

20. Grahn D, Wright BJ, Carnes B, Williamson FS, Fox C. Studies of Acute and Chronic Radiation Injury at the Biological and Medical Research Division, Argonne National Laboratory, 1970-1992: the JANUS Program Survival and Pathology 1995.

21. Haley B. Janus Tissue Archive Northwestern University 2010 [Available from: http://janus.northwestern.edu/janus2/index.php.

22. Haley B. Past and Future Work on Radiobiology Mega Studies: A Case Study at Argonne National Laboratory. 2011;100(6):613–21.

23. Birschwilks M, Schofield PN, Grosche B. The European Radiobiological Archives: online access to data from radiobiological experiments is available now. Health Phys. 2012;102(2):220.

24. Tanaka IB, 3rd, Tanaka S, Ichinohe K, Matsushita S, Matsumoto T, Otsu H, et al. Cause of death and neoplasia in mice continuously exposed to very low dose rates of gamma rays. Radiation research. 2007;167(4):417–37.

25. Cox DR. Regression Models and Life-Tables. Journal of the Royal Statistical Society. 1972;34(2):187–220.

26. Austin PC, Lee DS, Fine JP. Introduction to the Analysis of Survival Data in the Presence of Competing Risks. Circulation. 2016;133(6):601–9.

27. Zhang Z. Survival analysis in the presence of competing risks. Ann Transl Med. 2017;5(3):47.

28. Zander A, Paunesku T, Woloschak G. Radiation databases and archives - examples and comparisons. Int J Radiat Biol. 2019:1–12.

29. Wang Q, Paunesku T, Woloschak G. Tissue and data archives from irradiation experiments conducted at Argonne National Laboratory over a period of four decades. Radiat Environ Biophys. 2010;49(3):317–24.

30. Liu W, Haley B, Kwasny MJ, Li JJ, Grdina DJ, Paunesku T, et al. Comparing radiation toxicities across species: an examination of radiation effects in Mus musculus and Peromyscus leucopus. Int J Radiat Biol. 2013;89(6):391–400.

31. Paunesku D, Paunesku T, Wahl A, Kataoka Y, Murley J, Grdina DJ, et al. Incidence of tissue toxicities in gamma ray and fission neutron-exposed mice treated with Amifostine. Int J Radiat Biol. 2008;84(8):623–34.

32. Rich JT, Neely JG, Paniello RC, Voelker CC, Nussenbaum B, Wang EW. A practical guide to understanding Kaplan-Meier curves. Otolaryngol Head Neck Surg. 2010;143(3):331–6.

33. Goel MK, Khanna P, Kishore J. Understanding survival analysis: Kaplan-Meier estimate. Int J Ayurveda Res. 2010;1(4):274–8.

34. Therneau TM. A Package for Survival Analysis in S 2015 [version 2.38:[Available from: https://CRAN.R-project.org/package=survival.

35. Fine JP, Gray RJ. A Proportional Hazards Model for the Subdistribution of a Competing Risk. Journal of the American Statistical Association. 1999;94(446):496–509.

36. Gray B. Subdistribution Analysis of Competing Risks 2014 [Available from: https://CRAN.R-project.org/package=cmprsk.

37. Latouche A, Allignol A, Beyersmann J, Labopin M, Fine JP. A competing risks analysis should report results on all cause-specific hazards and cumulative incidence functions. J Clin Epidemiol. 2013;66(6):648–53.

38. Noordzij M, Leffondre K, van Stralen KJ, Zoccali C, Dekker FW, Jager KJ. When do we need competing risks methods for survival analysis in nephrology? Nephrol Dial Transplant. 2013;28(11):2670–7.

39. Liu W, Haley BM, Kwasny MJ, Li JJ, Grdina DJ, Paunesku T, et al. The effects of radiation and dose-fractionation on cancer and non-tumor disease development. Int J Environ Res Public Health. 2012;9(12):4688–703.

40. Braga-Tanaka I, 3rd, Tanaka S, Kohda A, Takai D, Nakamura S, Ono T, et al. Experimental studies on the biological effects of chronic low dose-rate radiation exposure in mice: overview of the studies at the Institute for Environmental Sciences. Int J Radiat Biol. 2018;94(5):423–33.

41. Grahn D. Acute Radiation Response of Mice from a Cross between Radiosensitive and Radioresistant Strains. Genetics. 1958;43(5):835–43.

42. Carnes BA, Gavrilova N, Grahn D. Pathology effects at radiation doses below those causing increased mortality. Radiation research. 2002;158(2):187–94.

43. Carnes BA, Grahn D, Thomson JF. Dose-response modeling of life shortening in a retrospective analysis of the combined data from the JANUS program at Argonne National Laboratory. Radiation research. 1989;119(1):39–56.

44. Grahn D, Lombard LS, Carnes BA. The comparative tumorigenic effects of fission neutrons and cobalt-60 gamma rays in the B6CF1 mouse. Radiation research. 1992;129(1):19–36.

45. Thomson JF, Grahn D. Life shortening in mice exposed to fission neutrons and gamma rays. VII. Effects of 60 once-weekly exposures. Radiation research. 1988;115(2):347–60.

46. Carnes BA, Grahn D, Hoel D. Mortality of atomic bomb survivors predicted from laboratory animals. Radiation research. 2003;160(2):159–67.

47. Hoel DG, Carnes BA. Cardiovascular effects of fission neutron or (60)Co gamma exposure in the B6CF1 mouse. Int J Radiat Biol. 2017;93(6):563–8.

48. Tuli JS, Smith JA, Morton DB. Stress measurements in mice after transportation. Lab Anim. 1995;29(2):132–8.

49. Gerdin AK, Igosheva N, Roberson LA, Ismail O, Karp N, Sanderson M, et al. Experimental and husbandry procedures as potential modifiers of the results of phenotyping tests. Physiol Behav. 2012;106(5):602–11.

50. Heidenreich WF, Carnes BA, Paretzke HG. Lung cancer risk in mice: analysis of fractionation effects and neutron RBE with a biologically motivated model. Radiation research. 2006;166(5):794–801.

51. Rivina L, Davoren MJ, Schiestl RH. Mouse models for radiation-induced cancers. Mutagenesis. 2016;31(5):491–509.

52. Ullrich RL, Jernigan MC, Adams LM. Induction of lung tumors in RFM mice after localized exposures to X rays or neutrons. Radiation research. 1979;80(3):464–73.

53. Yuhas JM, Walker AE. Exposure-response curve for radiation-induced lung tumors in the mouse. Radiation research. 1973;54(2):261–73.

54. Cahoon EK, Preston DL, Pierce DA, Grant E, Brenner AV, Mabuchi K, et al. Lung, Laryngeal and Other Respiratory Cancer Incidence among Japanese Atomic Bomb Survivors: An Updated Analysis from 1958 through 2009. Radiation research. 2017;187(5):538–48.

55. Egawa H, Furukawa K, Preston D, Funamoto S, Yonehara S, Matsuo T, et al. Radiation and smoking effects on lung cancer incidence by histological types among atomic bomb survivors. Radiation research. 2012;178(3):191–201.

